# Deletion of MOrpholino Binding Sites (DeMOBS) to Assess Specificity of Morphant Phenotypes

**DOI:** 10.1101/2020.01.12.903211

**Authors:** Carlee MacPherson Cunningham, Gianfranco Bellipanni, Raymond Habas, Darius Balciunas

## Abstract

Two complimentary approaches are widely used to study gene function in zebrafish: induction of genetic mutations, usually using targeted nucleases such as CRISPR/Cas9, and suppression of gene expression, typically using Morpholino oligomers. Neither method is perfect. Morpholinos (MOs) sometimes produce off-target or toxicity-related effects that can be mistaken for true phenotypes. Conversely, genetic mutants can be subject to compensation, or may fail to yield a null phenotype due to leakiness. When discrepancy between mutant and morpholino-induced (morphant) phenotypes is observed, experimental validation of such phenotypes becomes very labor intensive. We have developed a simple genetic method to differentiate between genuine morphant phenotypes and those produced due to off-target effects. We speculated that indels within 5’ untranslated regions would be unlikely to have a significant negative effect on gene expression. Mutations induced within a MO target site would result in a Morpholino-refractive allele thus suppressing true MO phenotypes whilst non-specific phenotypes would remain. We tested this hypothesis on one gene with an exclusively zygotic function, *tbx5a*, and one gene with strong maternal effect, *ctnnb2*. We found that indels within the Morpholino binding site are indeed able to suppress both zygotic and maternal morphant phenotypes. We also observed that the ability of such indels to suppress Morpholino phenotypes does depend on the size and the location of the deletion. Nonetheless, mutating the morpholino binding sites in both maternal and zygotic genes can ascertain the specificity of morphant phenotypes.

## Introduction

Methods used to analyze gene loss-of-function fall into two categories: “knockout”, which aims to inactivate the gene of interest by introducing mutations using techniques such as CRISPR/Cas9, and “knockdown”, which aims to abrogate the expression of the gene of interest by employing methods such as siRNA, CRISPRi or Morpholino oligomers (for a recent review, see (Housden et al., 2017)).

Morpholino oligomers are the most widely used antisense knockdown technology in the zebrafish (Ekker and Larson, 2001; Nasevicius and Ekker, 2000). They inhibit gene expression by blocking either translation or splicing. Translation-blocking MOs base pair with the mRNA at or upstream of the translation start site and prevent assembly of the 80S ribosome. They inhibit expression of both zygotic and maternally deposited mRNAs and can be used to phenocopy maternal-zygotic mutants. Splice-blocking MOs bind to pre-mRNA at either the splice acceptor or the splice donor site and prevent assembly of the spliceosome, thus abrogating expression of zygotic transcripts but having no effect on maternally deposited mature mRNAs. MO technology, however, uffers from a significant drawback: off-target effects. Some off-target effects caused by activation of the p53 pathway can be suppressed by co-injection of a standard MO targeting p53 (Robu et al., 2007). Nonetheless, presence of off-target effects necessitates thorough and labor-intensive validation of morphant phenotypes, including mRNA rescue and use of multiple MOs targeting the gene of interest (Eisen and Smith, 2008; Stainier et al., 2017).

It is not uncommon for knockdown- and knockout-based approaches to yield different results (Bachas et al., 2018; Evers et al., 2016; Luttrell et al., 2018; Morgens et al., 2016). Such discrepancies have also been observed in zebrafish (Joris et al., 2017; Kok et al., 2015; Law and Sargent, 2014; Novodvorsky et al., 2015; Rossi et al., 2015). Sometimes they can be attributed to built-in shortcomings of each approach. Small indels do not always result in complete loss-of-function: a variety of phenomena including splicing artifacts and translation initiation at a downstream AUGs and may lead to production of a functional protein, masking the null phenotype (Anderson et al., 2017; Lalonde et al., 2017; Smits et al., 2019). A further complication in the analysis of mutant phenotypes arises from the fact that for some genes, maternally-deposited mRNAs partly mask mutant phenotypes necessitating the use of maternal-zygotic mutants (Gritsman et al., 1999; Miller-Bertoglio et al., 1999). Additionally, some frameshift and nonsense mutants induce transcriptional compensation by closely related genes (El-Brolosy et al., 2019; Ma et al., 2019; Rossi et al., 2015). Deleting the whole coding sequence appears to be the best way to eliminate these possibilities. However, regulatory complexity of vertebrate genomes raises the possibility the observed phenotype may be caused deletion of intron-residing *cis*-regulatory elements for other genes (for an example, see (Zhou et al., 2009; Zuniga et al., 2012)).

With the notable exception of short upstream reading frames (Johnstone et al., 2016), 5’ UTRs appear to be sparse in significant regulatory features. We speculated that indel mutations within 5’ UTRs are unlikely to significantly impair the expression of the downstream gene (Burg et al., 2018). Introduced indels within a morpholino target site should reduce, if not entirely abolish, MO binding making the “mutant mRNA” partly or completely refractive to morpholino binding. We further hypothesized that since few genes are haploinsufficient, heterozygocity for such MO-refractive mutations would be sufficient to suppress specific morpholino phenotypes. Using *tbx5a* and *ctnnb2* as test loci, we demonstrate that deletions can be readily generated and used to test the specificity of both zygotic and maternal morphant phenotypes.

## Results and Discussion

### Partial suppression of *tbx5a* morphant phenotypes by the (−7) mutation in MO target site

Tbx5a mutants and morphants display absent or malformed pectoral fins and a linear heart (**Figure 1A**) (Ahn et al., 2002; Garrity et al., 2002; Grajevskaja et al., 2018; Ng et al., 2002). Since *tbx5a* mRNA is not contributed maternally, outcross of a parent heterozygous for a potentially MO-refractive mutation would produce a clutch of embryos where half would be genotypically wild type and susceptible to the MO, while the other half would be “mutant” and thus refractive to the MO. Susceptible and refractive embryos, present within a single clutch, would serve as controls for each other, eliminating experimental bias by excluding variables such as active MO concentration, injection volume or timing of the injection.

**Figure 1.**
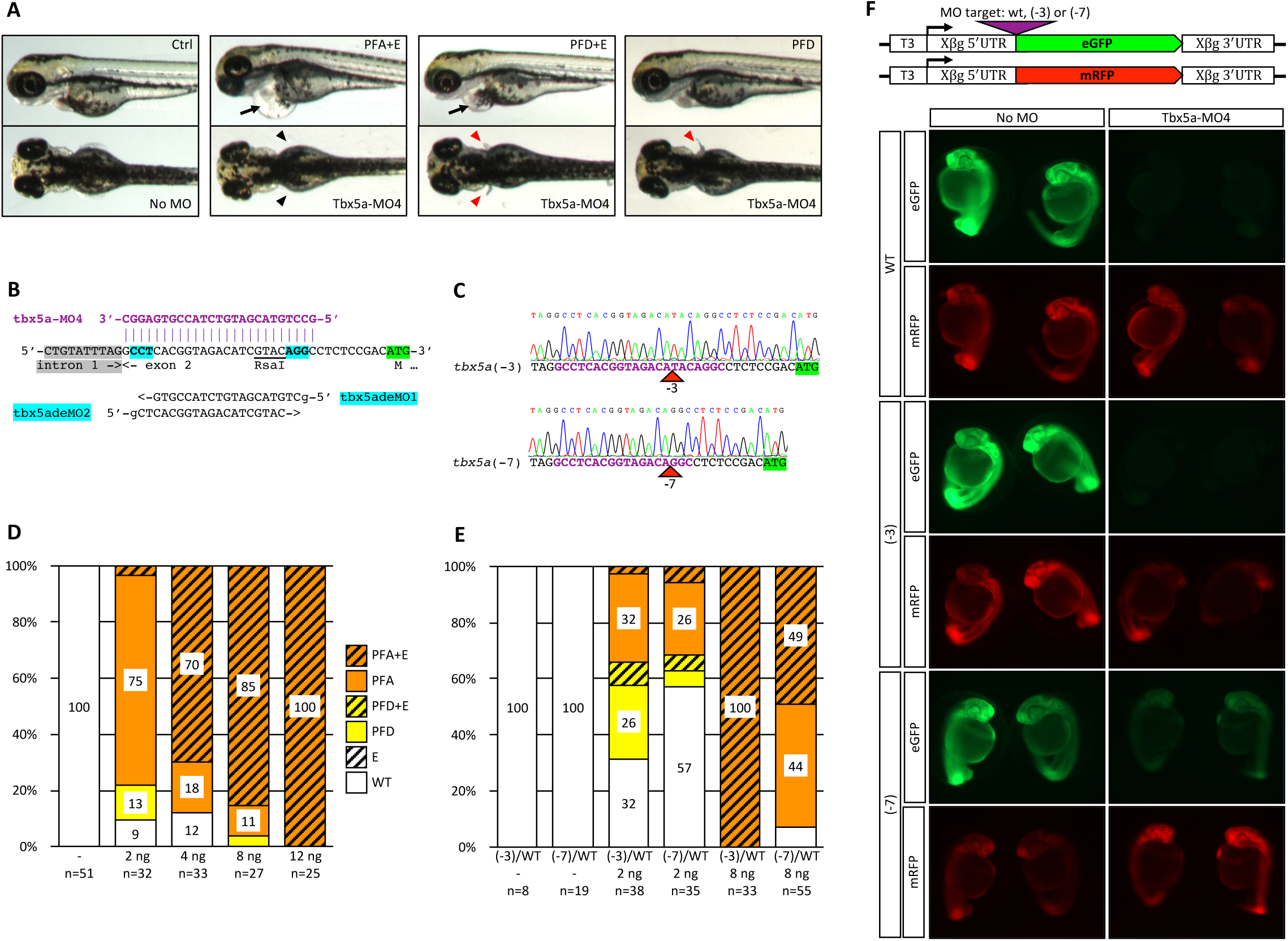
Partial rescue of Tbx5a-MO4 morphant phenotype by (−3) and (−7) binding site mutations. (A) Embryos injected with Tbx5a-MO4 display a range of *tbx5a* loss of function phenotypes including pectoral fin malformation or absence and severe cardiac edema. Black arrow denotes cardiac edema, black arrowheads denote pectoral fin loss, red arrowheads denote pectoral fin defects. (PFA+E, <u>p</u>ectoral fins <u>a</u>bsent with <u>e</u>dema, PFD+E, <u>p</u>ectoral fin <u>d</u>efect with <u>e</u>dema, PFD, <u>p</u>ectoral fin <u>d</u>efect only) (B) Two sgRNAs, tbx5adeMO1 and tbx5adeMO2 targeting the tbx5a-MO4 binding site (MO sequence shown above in purple, PAM sites are highlighted in magenta, coding sequence is highlighted in green. Tbx5adeMO2 overlaps a RsaI restriction enzyme site used for genotyping. (C) Sequence confirmation of the (−3) and (−7) deletion alleles. (D) Titration of Tbx5a-MO4 in wild type (TFL) embryos. Numbers indicate percentages of embryos displaying designated morphant phenotypes. WT is wild-type, E is edema only. (E) Suppression of cardiac and/or pectoral fin phenotypes by the (−3) and (−7) binding site deletions at different doses of MO. Adults heterozygous for either the (−3) or (−7) deletion were outcrossed and embryos were injected with either 2 ng or 8 ng of tbx5a-MO4. (F) Tbx5a-MO4 is able to at least partly block the translation of mRNAs containing (−3) and (−7) binding site mutations. *In vitro* transcribed mRNAs from each eGFP construct was injected along with mRFP mRNA as a control, and half of the mRNA-injected embryos were then injected with tbx5a-MO4. At 1dpf, embryos displaying similar levels of mRFP expression were photographed. T3, T3 transcription start site, Xβg 5’ UTR and Xβg 3’ UTR, Xenopus β-globin 5’ and 3’ untranslated regions, respectively.

Two *S. pyogenes* PAM sites are present within the 5’UTR sequence targeted by tbx5a-MO4 (Lu et al., 2008) (**Figure 1B**). We synthesized 19-base guide RNAs with a G nucleotide (lower case in **Figure 1B**) required for transcription initiation by the T7 RNA polymerase added, and injected them along with nCas9n mRNA as previously described (Burg et al., 2018; Burg et al., 2016). PCR fragments amplified on lysates from 20 pooled 3 day post fertilization (dpf) injected embryos were analyzed for sgRNA efficiency by TIDE (Brinkman et al., 2014) and Synthego ICE (Hsiau et al., 2019). Both analyses showed that tbx5adeMO2 sgRNA (∼30% by TIDE, ∼11% by ICE) was more efficient than tbx5adeMO1 sgRNA (∼17% by TIDE, 7% by ICE) (**Supplemental Figure 1A)**. We raised embryos injected with tbx5adeMO2 sgRNA and nCas9n and screened three F0 fish for germline transmission of indels using the T7 endonuclease assay (data not shown). One founder produced a high percentage of progeny with indels, and one F1 family was raised. Four out of seven genotyped adult F1s were found to be heterozygous for indels: two for a (−3) deletion and two for a (−7) deletion by Poly Peak Parser (Hill et al., 2014) analysis. F1s heterozygous for (−3) and (−7) deletions were incrossed. All embryos were phenotypically normal, indicating that these deletions do not significantly impair the expression of *tbx5a*. Sequence of the (−3) and (−7) deletions was confirmed on homozygous F2s (**Figure 1C**).

To determine the effective dose of Tbx5a-MO4, we injected 2, 4, 8 and 12 ng of the morpholino into one-cell zebrafish embryos (**Figure 1D**). The lowest dose of the MO resulted in >90% of embryos displaying pectoral fin defects. In contrast, a much higher 8 ng dose of the MO was needed to elicit severe cardiac defects (>90% edema). Notably, in humans suffering from Holt-Oram syndrome caused by mutations in Tbx5, forelimb defects are also more penetrant and severe than cardiac defects (Basson et al., 1997; Basson et al., 1994; Smith et al., 1979).

We outcrossed F1 fish heterozygous for the (−3) and (−7) deletions and injected embryos with 2 ng or 8 ng Tbx5a-MO4. In each cross, approximately 50% of embryos were expected to be genetically wild type and therefore display morphant phenotypes, while the other 50% were expected to inherit the corresponding deletion, leading to either partial or complete suppression of MO phenotypes. At the low 2 ng MO dose, 12/38 (32%) of embryos from the (−3) heterozygote outcross were phenotypically wild type, whilst 10/38 (26%) showed the milder pectoral fin defect phenotype, indicating full or partial rescue in approximately 50% of the progeny as expected. Among embryos form the (−7) heterozygote outcross, 20/35 (57%) showed full rescue of both the cardiac edema and pectoral fin loss phenotype (**Figure 1E**). Embryos from both outcrosses were grouped by phenotype and genotyped. In the (−3) outcross, genotyping revealed that 6/8 (75%) embryos displaying pectoral fin loss or defects were wild-type, and 7/8 (88%) phenotypically wild-type embryos were heterozygous for the (−3) deletion (*P* = .015). In the (−7) outcross, genotyping revealed that 8/8 (100%) embryos displaying pectoral fin loss or defects were wild-type, and 7/8 (88%) phenotypically wild-type embryos were heterozygous for the (−7) deletion (*P* = .002). These results confirmed that at a low dose of the morpholino, the (−3) allele can largely suppress the morphant phenotype while the (−7) allele shows complete rescue.

At the high 8 ng MO dose, all MO-injected embryos from the (−3) outcrosses displayed identical morphant phenotypes (**Figure1E**). In contrast, approximately 50% of embryos from the (−7) outcross were completely rescued from the cardiac edema phenotype, but not the pectoral fin defect (**Figure 1E**). Embryos from the (−7) outcross were grouped by phenotype and subsequently genotyped. Genotyping revealed that 14/15 (93%) of individuals with cardiac edema and loss of pectoral fins were wild-type and 13/16 (81%) of individuals with no cardiac edema were heterozygous for the (−7) allele (*P*=3.0E-05) (data not shown). These results indicate rescue of cardiac edema morphant phenotype, but not the pectoral fin phenotype, by heterozygocity for the (−7) allele at the high dose of the Morpholino.

Dose-, phenotype- and deletion size-dependent rescue of morphant phenotypes prompted us to hypothesize that (−3) and (−7) deletions were insufficient to make mRNAs entirely refractive to the morpholino. We cloned wild type, (−3) and (−7) target sites into the pT3TS *in vitro* transcription vector (Hyatt and Ekker, 1999) ahead of eGFP coding sequence. *In vitro* transcribed mRNAs were injected into embryos along with pT3TS:mRFP mRNA as a control not affected by the MO. Half of the mRNA-injected embryos were then injected with 8 ng of Tbx5a-MO4. Embryos were scored for RFP and GFP fluorescence and photographed at 1 dpf (**Figure 1F**). We found that Tbx5a-MO4 was able to almost entirely block translation of mRNAs containing wild type and (−3) target sites. Translation of mRNA containing the (−7) target site was also reduced significantly (**Figure 1F)**. These findings raise significant concerns about MO specificity. The (−3) and (−7) deletions preserved a 16-nucleotide and 15-nucleotide identity to the target site at the 3’ end of the MO (toward the 5’ end of the mRNA), respectively. Our data therefore indicate that a 3’ stretch of identity as short 15-16 nucleotides may be sufficient for a MO to impede translation at a high dose, and suggests that using MOs shorter than the current 25 nucleotide standard may lead to higher specificity.

### Suppression of the *b-catenin* morphant phenotype by maternal contribution of (−4) binding site mutant *ctnnb2* mRNA

We speculated that for maternally contributed genes, a female heterozygous for a MO-refractive allele would produce embryos which would be refractive to the maternal-zygotic phenotype. To test this hypothesis, we selected beta-catenin genes coding for an essential component of the Wnt signaling pathway. Co-injection of translation-blocking MOs targeting the duplicated *ctnnb1* and *ctnnb2* mRNAs results in complete loss of ventral cell fates and a phenotype named *ciuffo* (Bellipanni et al., 2006). Partial sequencing of *ctnnb2* loci in the TLF genetic background revealed presence of a single nucleotide polymorphism within the ctnnb2-MO1 binding site. Co-injection of ctnnb1-MO2 and ctnnb2-MO1 into TLF embryos at concentrations described previously (Bellipanni et al., 2006) resulted in nearly 100% penetrant *ciuffo* phenotype (data not shown), indicating the polymorphism alone does not appreciably reduce morpholino activity.

Two sgRNAs targeting PAM sequences within the ctnnb2-MO1 binding site were designed and tested (**Figure 2A**). Only ctnnb2deMO1 had detectable activity by both TIDE (∼17%) and Synthego ICE (∼5%) analysis (**Supplementary Figure 2**). Three adult F0 fish injected with ctnnb2deMO1 sgRNA and nCas9n mRNA were tested for germline transmission of indels, leading to establishment of one F1 family. Fourteen F1 fish were tested for loss of Bpu10I restriction enzyme site, and seven were found to be heterozygous for indels. PCR fragments from five F1s were sequenced, and four were found to be heterozygous for a (−4) deletion (**Figure 2B**). A male heterozygous for the (−4) deletion was outcrossed to establish an F2 family.

**Figure 2.**
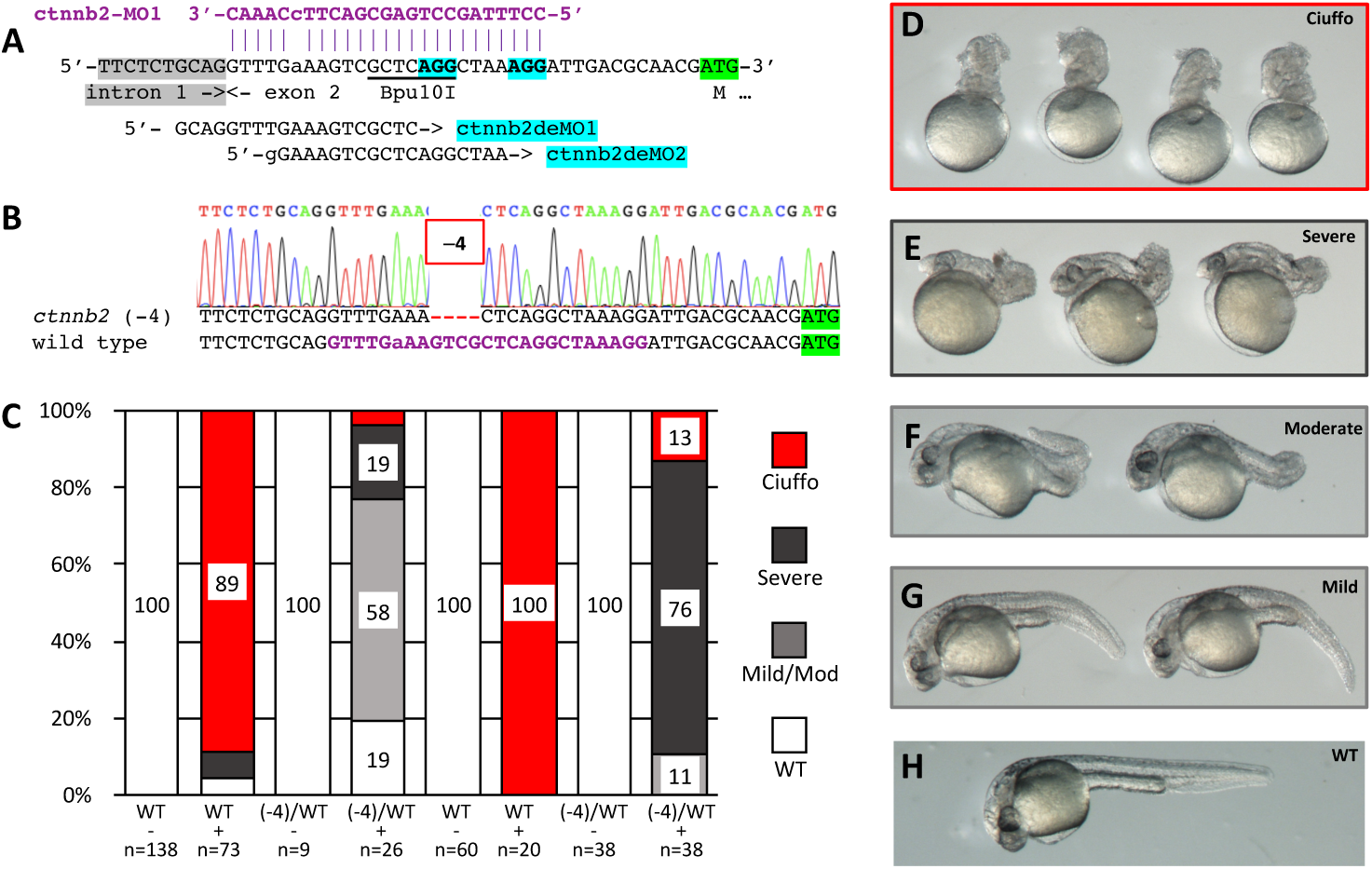
Suppression of a maternal morphant phenotype by DeMOBS. (A) Two sgRNAs, ctnnb2deMO1 and ctnnb2deMO2 target ctnnb2-MO1 binding site. ctnnb2deMO1 overlaps a Bpu10I restriction enzyme site used for genotyping. A single nucleotide polymorphism resulting in a single base mismatch between the MO and the target sequence is shown in lower case. (B) Confirmation of the (−4) allele by sequence analysis. (C) Co-injection ctnnb2-MO1 and ctnnb1-MO2 into embryos obtained from four female siblings (two wild type and two heterozygous for the deletion) in blind experiments. (D) *ciuffo* morphant phenotype resulting from injection of ctnnb1-MO2 and ctnnb2-MO1. (E-H) Residual phenotypes observed in MO-injected embryos from a female heterozygous for the MO-refractive (−4) mutation.

To avoid experimental bias, we performed a blind experiment to test for suppression of *ciuffo* phenotype. From a single F2 family, we identified two adult females heterozygous for the deletion and two wild type siblings. Fish were coded A, B, C and D, and embryos obtained from outcrosses were injected with a mixture of ctnnb1-MO2 and ctnnb2-MO1. MO injection lead to high penetrance (89% and 100%) of *ciuffo* phenotypes in the progeny of wild type fish, while presence of one (−4) allele in heterozygotes almost completely suppressed the *ciuffo* phenotype (4% and 13%, **Figure 2C**). Milder phenotypes were still observed in a subset of embryos, which could reflect zygotic requirement for *ctnnb1* and/or *ctnnb2* (Valenti et al., 2015; Varga et al., 2007), or off-target effects due to a high cumulative dose of the two MOs. Nonetheless, the observation that females heterozygous for a (−4) deletion in combination with a serendipitous single nucleotide polymorphism produce embryos which are nearly 100% suppressed for the *ciuffo* phenotype indicates that out method can be used to ascertain morphant phenotypes of genes with strong maternal contribution of mRNA.

Our data clearly demonstrates that indels within MO binding sites can be readily generated and used to test the specificity of both zygotic and maternal morphant phenotypes. The ability to induce deletions within MO binding sites using CRISPR/Cas9 relies on the presence of a PAM site within the MO binding site, preferably close to the 5’ end of the target site. While the mutagenesis method employed by us is likely feasible for the majority of MOs targeting 5’ UTRs, there will inevitably be a subset where PAM sites will be absent or located closer to the middle or the 3’ of the target site. In such cases, oligonucleotide-mediated repair of double strand breaks can be used to engineer desired mutations (Bedell et al., 2012; Burg et al., 2018; Burg et al., 2016; Dong et al., 2014; Gagnon et al., 2014; Gibb et al., 2018; Hruscha et al., 2013; Prykhozhij et al., 2018).

## Materials and Methods

### CRISPR/Cas9 mutagenesis

Guide RNAs were produced as previously described (Burg et al., 2018; Burg et al., 2016) using DR274 (Hwang et al., 2013) as the template and diluted to ∼60 ng/ μL. Immediately prior to injection, 8 uL of diluted sgRNA was mixed with 2 μL aliquot of 150 ng/ μL nCas9n mRNA (Jao et al., 2013) to the final volume of 10 μL.

### Plasmid construction and mRNA synthesis

Details of plasmid construction are available upon request. eGFP-containing pT3TS (Hyatt and Ekker, 1999) vectors (pCMC23 (wt MO binding site), pCMC24 (−3) and pCMC25 (−7)) were linearized using XbaI restriction enzyme. Template for the synthesis of mRFP mRNA was amplified by PCR using M13F/M13R primer pair on pT3TS:mRFP (pDB935). Templates were transcribed using T3 mMessage mMachine kit and mRNAs were purified using Qiagen RNeasy MinElute kit. mRNAs were diluted so that the standard 3 nL injection volume would contain 50 ng of tbx5a-eGFP mRNA and 100 ng of mRFP mRNA.

### Microinjection

Microinjection volumes were calibrated to 3 nL as previously described, and all microinjections were performed into the yolks of 1-cell zebrafish embryos as described (Balciuniene and Balciunas, 2013).

## Figure Legends

**Supp. Figure 1.**
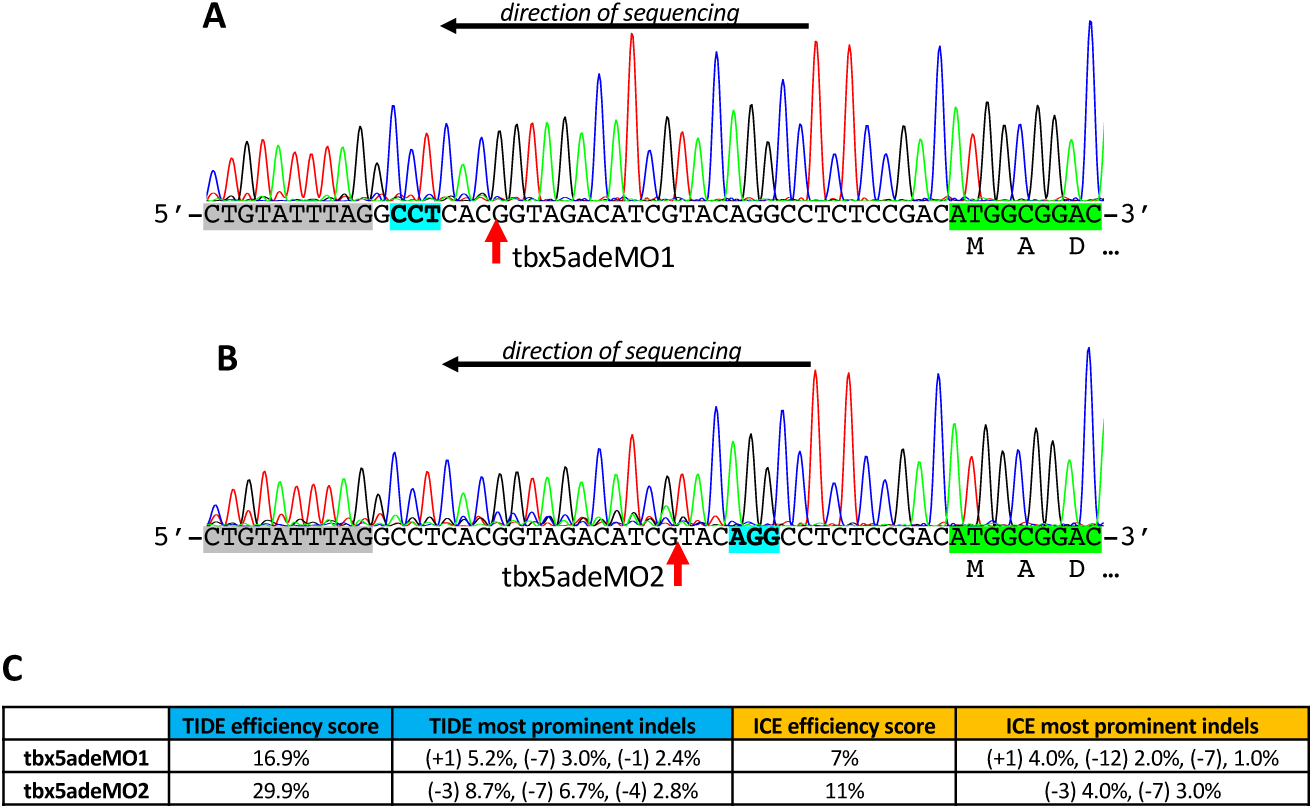
Testing the efficiency tbx5adeMO1 or tbx5adeMO2 sgRNAs by sequencing. Targeted locus was amplified with primers tbx5ain1-F2 and tbx5aex2-R2, using lysate of 20 pooled embryos as the template. PCR fragments were sequenced using tbx5aex2-R2. (A) Sequencing of embryos injected with tbx5adeMO1. Please note that reverse complement of sequencing chromatogram is provided to correspond to Figure 1B. The expected location of the double strand break is indicated by the red arrow. (B) Sequencing of embryos injected with tbx5adeMO1. (C) Efficiency of the two guides and most prevalent deletions as estimated by TIDE (https://tide.deskgen.com/) and Synthego ICE (https://ice.synthego.com/#/).

**Supp. Figure 2.**
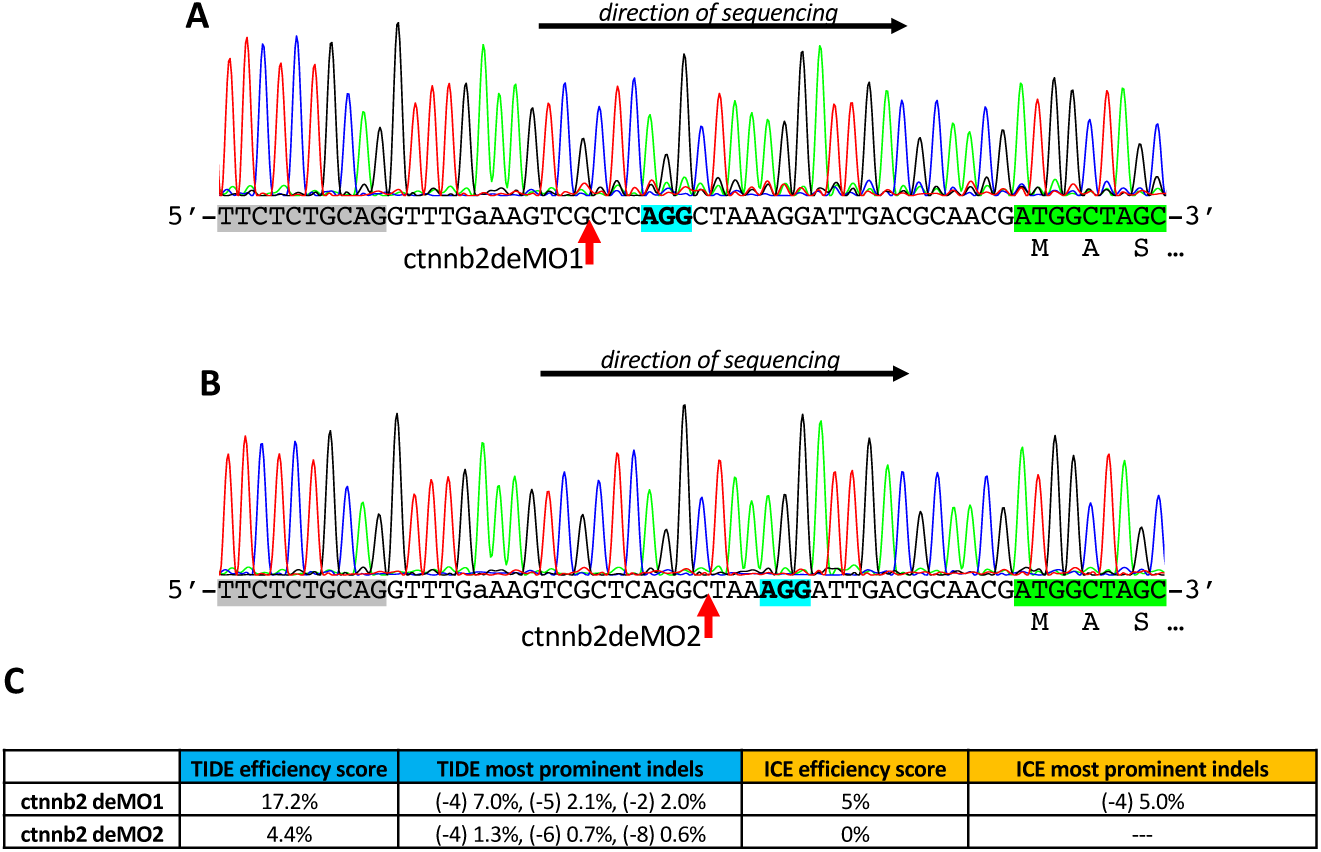
Testing the efficiency Testing the efficiency ctnnb2deMO1 or ctnnb2deMO2 guide RNAs by sequencing. Targeted locus was amplified with primers ctnnb2-F1 and ctnnb2-R1, using lysate of 20 pooled embryos as the template, and the obtained PCR fragments were sequenced using ctnnb2-F1. (A) Sequencing of embryos injected with ctnnb2deMO1. The expected location of the double strand break is indicated by the red arrow. Sequencing of embryos injected with ctnnb2deMO2. (C) Efficiency of the two guides and most prevalent deletions as estimated by TIDE (https://tide.deskgen.com/) and Synthego ICE (https://ice.synthego.com/#/).

**Supplementary Table 1.**
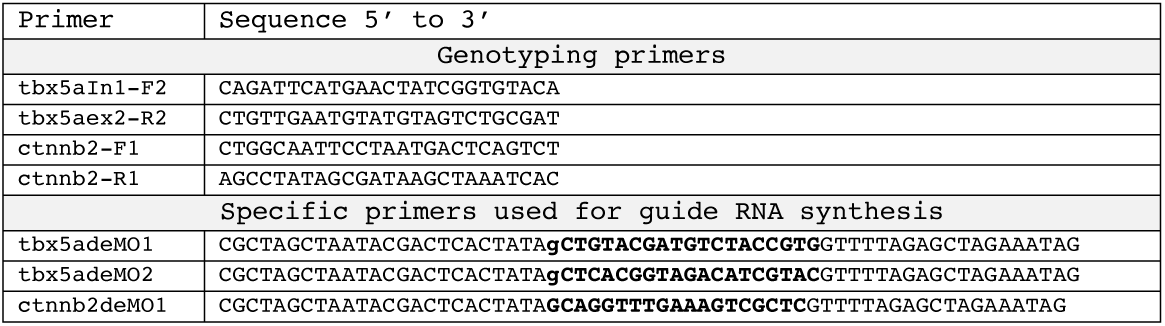
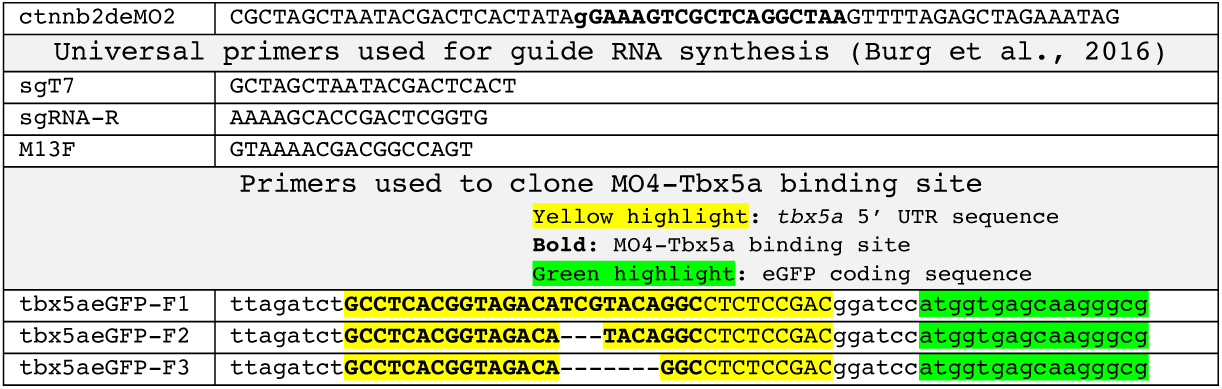
Primer sequences.

**Supplementary Table 2.**
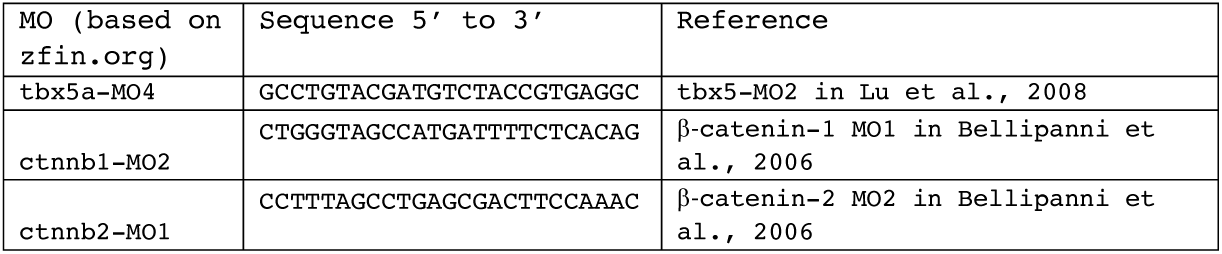
Sequences of Morpholino oligomers.

